# Cryo-EM structure and biochemical characterization of a BRAF/CRAF heterodimer: Negative charge in the NtA motif is not required for RAF activation

**DOI:** 10.64898/2026.05.11.724350

**Authors:** Byung Hak Ha, Emre Tkacik, Dimitrios Gazgalis, Hyelim Kang, Dong Man Jang, Sayan Chakraborty, Hyesung Jeon, Michael J. Eck

## Abstract

Upon RAS-driven membrane recruitment, RAF kinases ARAF, BRAF and CRAF are activated via formation of homo- or hetero-dimers to initiate signaling through the MAP kinase cascade. Although RAF heterodimers are important for both physiologic and oncogenic signaling, they have been little studied at a structural and biochemical level. Here we report the preparation, biochemical characterization, and the cryo-EM structure of a 14-3-3-bound BRAF/CRAF heterodimer complex. The heterodimer exhibited kinetic parameters and sensitivity to a panel of twelve structurally diverse RAF inhibitors that were closely similar to, or intermediate between, those of BRAF and CRAF homodimers. Cryo-EM structures of the heterodimer with and without MEK1 revealed an overall organization essentially identical to that of RAF homodimers, but with an asymmetric interaction in the MEK1-bound structure in which the BRAF N-terminal acidic (NtA) motif extends across the dimer interface to engage the CRAF RKTR motif. Mutagenesis of this interface unexpectedly revealed that replacing the acidic NtA sequence with a basic RARA sequence yields highly active RAF homodimers and heterodimers, demonstrating that negative charge in the NtA motif is not required for activity. Collectively, our findings suggest that the charge state of the NtA motif influences RAF activity through effects on local backbone dynamics and the stability of the inactive kinase conformation, rather than via stereospecific recognition across the dimer interface.

## Introduction

RAF-family kinases are a key link in the RAS/MAP kinase signal transduction cascade.^1^ The three RAF kinase isoforms ARAF, BRAF, and CRAF are activated by RAS-driven recruitment to the membrane which induces their dimerization or heterodimerization. Once active, RAFs phosphorylate their only substrates, the kinases MEK1 and MEK2. Structural studies of RAF complexes over the past several years have shown how RAFs are maintained in an autoinhibited state as a complex with MEK and a 14-3-3 dimer and how this complex reorganizes upon activation.^2,3^ In the autoinhibited state, the dimeric 14-3-3 subunit binds two serine phosphorylated sites that flank the RAF kinase domain. Together with stabilizing intramolecular interactions of the N-terminal regulatory RAS-binding and cysteine-rich domains, the 14-3-3 sterically blocks the dimer interface of the RAF kinase domain, ensuring that it remains monomeric and inactive. Membrane recruitments releases these inhibitory interactions, and the 14-3-3 rearranges to bind the C-terminal phosphoserine site in two RAFs, driving formation of an active, side by side dimer of the RAF kinase domains.

This stimulus-dependent activation mechanism is subverted by oncogenic mutations. The most common is the BRAF V600E point mutation that is a common cause of melanoma and papillary thyroid cancers, but also occurs in lung, colorectal and other cancers.^4^ This mutation lies in the activation loop of the kinase and allows BRAF to become active as a monomer.^5^ Other oncogenic activation mechanisms include chromosomal translocations that produce constitutively active RAF dimers due to truncation of the N-terminal regulatory domains and point mutations in the N-terminal 14-3-3 binding site that relieve autoinhibition.^6-8^ RAF inhibitors are important cancer therapeutics, in particular for cancers driven by BRAF V600E.^9^ As direct effectors of RAS, RAFs are also a prominent therapeutic target in RAS-altered cancers.^10^

Although structures are available for autoinhibited RAFs^11-13^ and for active homodimers of each of the three RAF isoforms^11,12,14-18^, RAF heterodimers have not been well-characterized at a structural or biochemical level. Furthermore, while overall regulatory mechanisms are conserved across RAFs, there are isoform-specific features that are not well-understood.^1,19^ In particular, ARAF and CRAF are thought to require phosphorylation of a tyrosine residue just N-terminal to their kinase domains for full catalytic activation.^20-23^ This tyrosine lies in a site known as the N-terminal acidic motif (NtA), and has the sequence SGYY in ARAF and SSYY in CRAF.^24^ The corresponding sequence in BRAF is SSDD, and one or both of the serine residues in this site are constitutively phosphorylated. The NtA motifs of ARAF and CRAF also undergo serine phosphorylation, although it is unclear whether this is constitutive or occurs only upon activation. Acidic substitutions of the tyrosine and serine residues in the ARAF and CRAF NtA motifs are highly activating (for example mutating SSYY to DDEE or to the SSDD sequence found in BRAF), and the evident requirement for negative charge in this region – whether via acidic mutation or phosphorylation – gave rise to the “N-terminal acidic” designation.^20,25-27^ Curiously, this segment has not been visualized in prior structures of RAFs in either autoinhibited or active states, and the mechanism by which it affects catalytic activity remains unclear.

In this study, we describe preparation, characterization, and structural analysis of a 14-3-3-bound BRAF/CRAF heterodimer. The BRAF/CRAF heterodimer complex was modestly more active than similarly prepared BRAF and CRAF homodimer complexes, and it exhibited Michaelis constants for substrates ATP and MEK1 that were closely similar to those of the homodimers. Inhibitor titrations with 12 structurally diverse inhibitors revealed sensitivities that were in the range observed for homodimers. Cryo-EM structures of this BRAF/CRAF heterodimer, with and without MEK1 bound to the BRAF protomer, reveal an overall heterodimer organization that is essentially the same as that observed for BRAF and CRAF homodimers bound to 14-3-3. However, in the MEK1-bound structure, we observe an asymmetric interaction of the BRAF NtA motif, which extends across the dimer interface to contact CRAF. Mutagenesis of this segment and of the interacting basic residues in CRAF yielded an entirely unexpected result – substitution of the NtA motif in BRAF and CRAF with a basic sequence (RARA) yields highly active RAF homodimers and heterodimers. Together with prior studies, our findings suggest that stereospecific interactions of this segment are not required for its effect on catalytic activity. Rather, the charge state of this segment may influence its dynamics and that of an immediately adjacent conserved tryptophan residue to alter the stability of the inactive, αC-helix-out conformation of the kinase.

## Results

### Preparation, characterization, and inhibitor sensitivity of BRAF/CRAF heterodimers

We co-expressed C-terminal constructs of BRAF and CRAF that spanned their kinase domains and C-terminal 14-3-3 binding sites using the baculovirus/insect cell expression system. Differential tagging of the two constructs (BRAF with a Strep-tag and CRAF with His_6_-tag) allowed isolation of BRAF/CRAF heterodimers by sequential affinity chromatography. The heterodimer co-purified with an insect-cell derived 14-3-3 dimer (Figure 1A). As in our prior work with CRAF, we incorporated the activating “SSDD” substitution for the native “SSYY” sequence of the CRAF NtA motif.^28^

**Figure 1.**
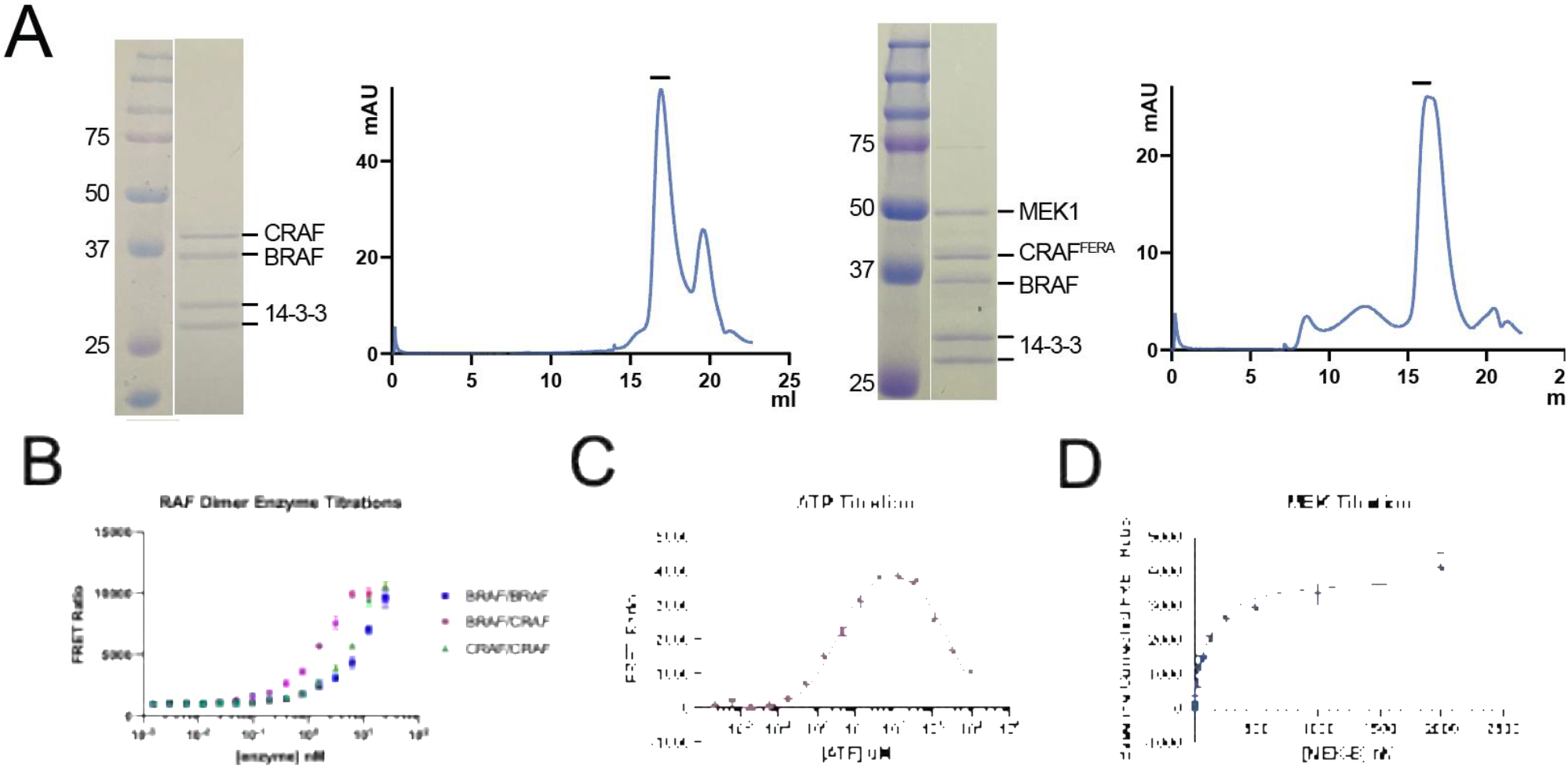
BRAF/CRAF heterodimer purification and activity. A, Coomassie-stained SDS-PAGE gels and corresponding size-exclusion chromatography profiles of purified BRAF/CRAF/14-3-3 (left) and BRAF/CRAF^FERA^/MEK1/14-3-3 (right) complexes. The CRAF employed here contains the SSDD NtA substitution and CRAF^FERA^ denotes CRAF containing both the SSDD substitution and the F559E/R563A mutations that prevent binding of MEK1 to CRAF. The bars above the SEC traces indicate fractions used for cryo-EM studies. B, Comparison of activity of 14-3-3-bound BRAF and CRAF homodimers and heterodimers assayed by TR-FRET. Signal ratio at 665/620 nm is plotted as a function of RAF dimer concentration. C, K_m[ATP_] determination for the BRAF/CRAF heterodimer (representative experiment from n=3). D, K_m[MEK]_ determination for the BRAF/CRAF heterodimer (representative experiment from n=3). The ATP titration was fit with a substrate inhibition model in GraphPad Prism to determine K_m[ATP]_ and K_i[ATP]_ values. The MEK1 titration was fit with the Michaelis-Menten equation in GraphPad Prism to determine K_m[MEK]_ values.

In our TR-FRET kinase assay, the purified BRAF/CRAF heterodimer was modestly more active than 14-3-3-bound BRAF and CRAF homodimers (Figure 1B). The Michaelis constant for ATP (K_m[ATP]_ = 2.3 µM, Figure 1C) was closely similar to values we previously reported for BRAF (1.9 µM) and CRAF (0.7 µM) homodimers.^28^ Similarly, affinity for substrate MEK1 (K_m[MEK]_ = 133 nM, Figure 1D) was also in line with values we have measured for BRAF and CRAF (30 to 150 nM).^28^ As with BRAF and CRAF homodimers, the heterodimer exhibits clear substrate inhibition at high ATP concentrations, with an apparent K_i_ value of approximately 3 mM (Figure 1C). This effect is thought to arise from binding of ATP to the canonical ATP-site, where it promotes an inactive conformation of the kinase ^11,15^.

We have previously examined the sensitivity of all three RAF isoforms to a panel of structurally diverse RAF inhibitors^28^, but the drug sensitivity of RAF heterodimers has received little direct study. Considering the role of RAF heterodimers in oncogenic signaling ^24,29^, we were interested to compare the inhibitor sensitivity of the heterodimer with that of homodimers. We profiled a panel of 12 diverse RAF inhibitors against BRAF homodimers, CRAF homodimers, and BRAF/CRAF heterodimers *in vitro*, using the same TR-FRET assay employed above. The IC_50_ and Hill Slope (nH) values of each enzyme against each inhibitor are reported in Table 1 and corresponding dose-response curves are shown in Supplementary Figure 1.

**Table 1.**
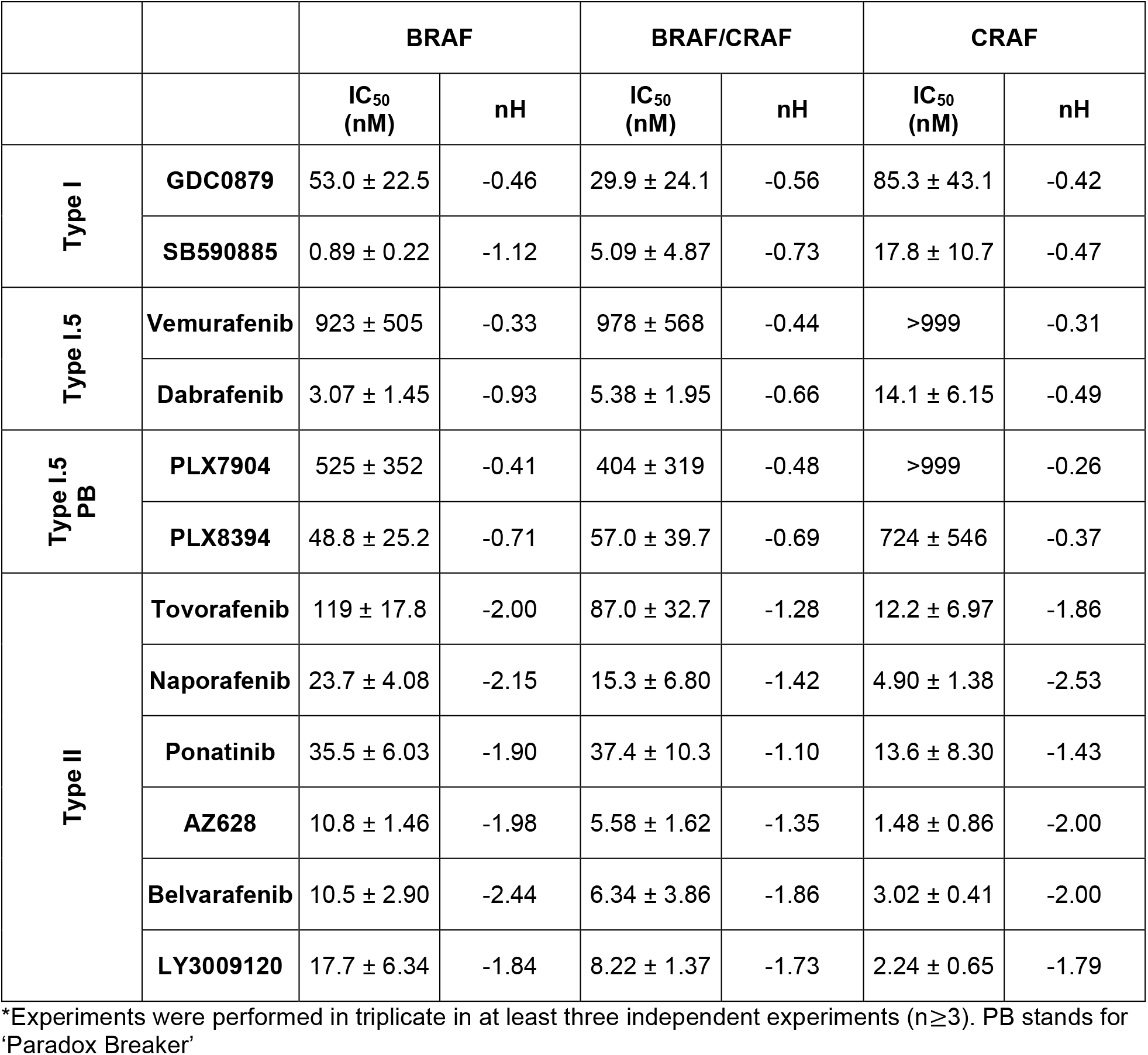
Enzymatic IC_50_ values for BRAF, CRAF, and BRAF/CRAF dimers.*

In general, potencies against BRAF/CRAF heterodimers were similar to or intermediate between those observed for the homodimers. This trend was evident across inhibitor classes. Between type I kinase inhibitors GDC0879 and SB590885, which bind the active *α*C-helix in, DFG-in conformation of the kinase, SB590885 was the more potent heterodimer inhibitor (IC_50_ = 5 nM). Type I.5 inhibitors bind the αC-helix-out conformation of the kinase and while they are very potent against the BRAF V600E mutant, which is active as a monomer ^5^, they are typically weak inhibitors of RAF dimers because dimerization promotes the active, αC-helix-in state. Dabrafenib was the exception among the four type I.5 inhibitors tested, with an IC_50_ of ~5 nM on the BRAF/CRAF heterodimer.Type II inhibitors bind a DFG-out conformation of the kinase and although they are often referred to as “pan-RAF” inhibitors, we previously showed that as a class they are most potent on CRAF and exhibit relative sparing of ARAF^28^. For the six type II inhibitors tested here, potencies against the heterodimer were within the range we observed for inhibition of BRAF and CRAF homodimers, but in general tracked more closely with potencies against BRAF homodimers (Supplementary Figure 1). As we previously reported, type II inhibitors inhibit BRAF and CRAF homodimers with apparent positive cooperativity (Hill slopes less than −1). This effect was less prominent with inhibition of the heterodimer (Table 1).

### Cryo-EM structures of BRAF/CRAF heterodimers

Co-expression of RAFs with MEK yields a higher-molecular weight complex that is in principle more amenable to single particle reconstruction. However, prior structures of MEK-bound RAF dimers have been C2-symmetric^11,16^, leading us to worry that the density corresponding to the closely similar BRAF and CRAF kinase domains would be averaged in single particle reconstructions. To obviate this concern, we introduced mutations in the MEK-binding interface of the CRAF kinase domain to disrupt its interaction with MEK1 (Phe559Glu/Arg563Ala, denoted CRAF^FERA^, Supplementary Figure 2A). We reasoned that the resulting asymmetry would prevent averaging of the similar kinase domains and also unambiguously mark BRAF as the MEK1-bound protomer. When prepared as a 14-3-3-bound homodimer, CRAF^FERA^ was not able to phosphorylate MEK1 and did not bind MEK1 in a pulldown assay (Supplementary Figure 2B,C). By co-expressing CRAF^FERA^ with BRAF and a MEK1 S218A/S222A mutant that is not phosphorylated by RAF, we obtained a BRAF/CRAF^FERA^ heterodimer that purified as a complex with MEK1 and 14-3-3 as expected (Figure 1A, right panels).

Cryo-EM imaging of the BRAF/CRAF/MEK1/14-3-3 preparation allowed single-particle reconstruction of two distinct complexes from the same image set; a 14-3-3-bound BRAF/CRAF heterodimer, and a similar complex but with MEK1 bound to the BRAF protomer (Figure 2A,B, Supplementary Figure 3, Supplementary Table 1). Counter to our expectations, reconstruction of the BRAF/CRAF/14-3-3 complex reached a higher resolution (3.3 Å) than did that of the larger complex containing MEK1 (3.9Å).

**Figure 2.**
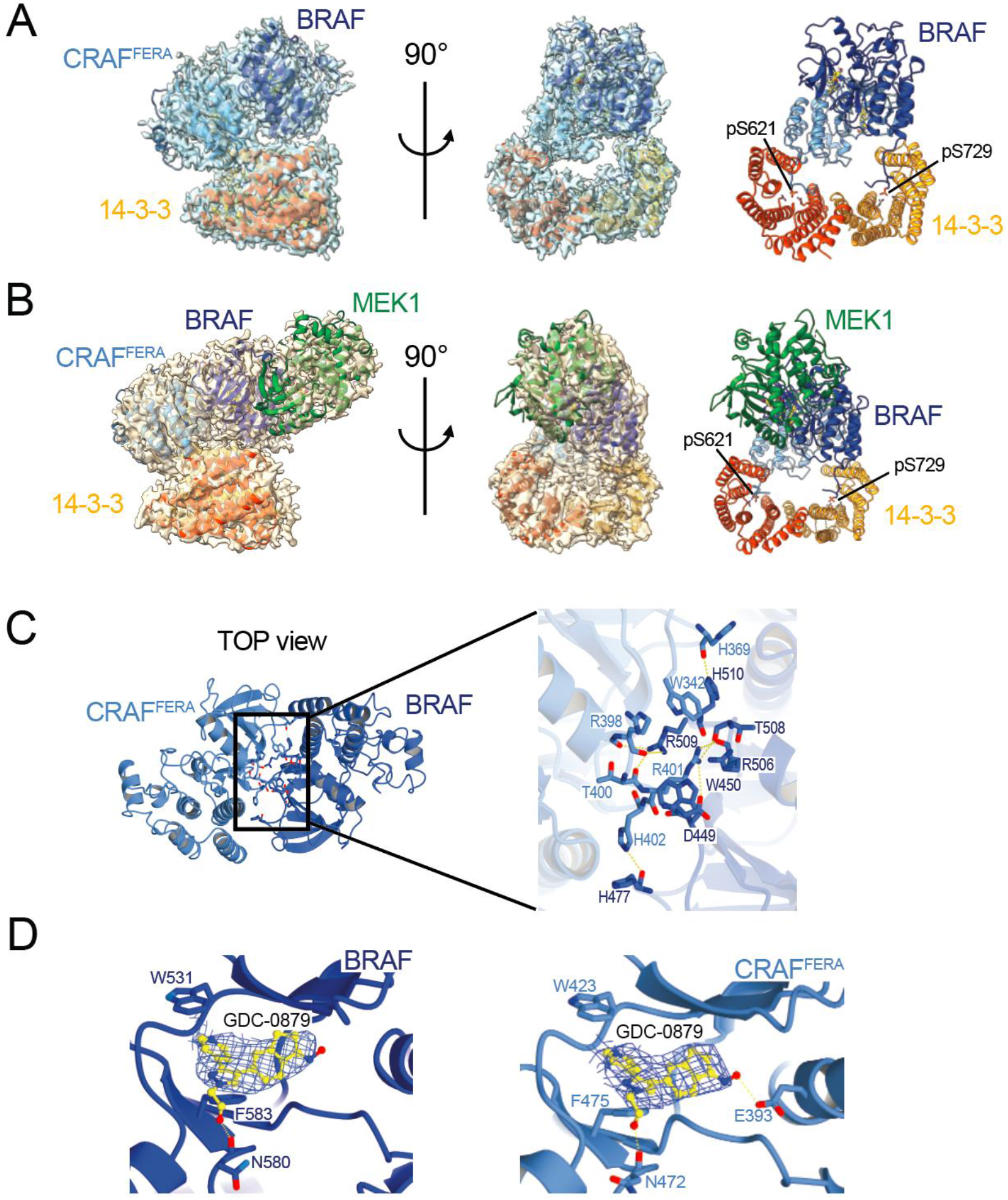
Cryo-EM structures of BRAF/CRAF heterodimers. A, BRAF/CRAF^FERA^/14-3-3 structure with cryo-EM map and orthogonal views of the complex. BRAF is shown in blue, CRAF^FERA^ in light blue, and the 14-3-3 dimer in orange and red-orange. B, Overall structure of the BRAF/CRAF^FERA^/MEK1/14-3-3 complex with map and orthogonal views of complex. The complex is colored as in A with MEK1 shown in green. Owing to the relatively weaker density for portions of MEK1, it was built based on a reference model (PDB ID 6PP9). C, Top view of BRAF/CRAF^FERA^ kinase dimer interface. Selected residues in the dimer interface are shown in stick form, with CRAF residues labeled in light blue and BRAF residues in dark blue. D, Cryo-EM density contoured in the region of inhibitor GDC-0879 bound to BRAF (left panel) and CRAF (right panel).

The BRAF/CRAF/14-3-3 reconstruction revealed an overall organization closely similar to that seen in 14-3-3-bound BRAF homodimers. The BRAF and CRAF kinase domains form the characteristic side by side dimer interaction that results in RAF activation, and their C-terminal tails are engaged by a 14-3-3 dimer (Figure 2A). While both the 14-3-3 and kinase domain dimers are pseudo 2-fold symmetric, their axes are not aligned and the overall structure is highly asymmetric. The kinase dimer tilts such that the CRAF kinase domain bridges across the 14-3-3 dimer, making contacts via both its N- and C-lobes, while the BRAF protomer makes contact with the 14-3-3 only via its C-terminal tail and immediately adjacent residues. These contacts are analogous to those seen in prior structures of 14-3-3-bound BRAF homodimers and the present structure superimposes on the BRAF heterodimer structure (PDB entry 6UAN^14^) with an overall RMSD of 1.65Å across 1007 aligned Cα residues (Supplementary Figure 4A). We have assigned the density corresponding to CRAF versus BRAF by careful examination of the cryo-EM map at sites where the corresponding amino acid sidechains differ in size (Supplementary Figure 4B).This assignment is consistent with that in the MEK-bound structure described below, where the upward-facing BRAF protomer associates with MEK1 (Figure 2B). However, we cannot exclude the possibility that there is some averaging of particles in which BRAF and CRAF assume the opposite positions. As evidenced by the BRAF homodimer structures, BRAF can make essentially the same 14-3-3 contacts^14^. The purified complex contains both the 14-3-3ε and 14-3-3ζ isoforms and we have modeled the protomer that binds the BRAF tail as 14-3-3ε and that CRAF-binding protomer as 14-3-3ζ. However, as in prior structures their positions appear to be degenerate. This is as expected considering their high sequence identity (~80 percent) and that they form both homo- and hetero-dimers.

The kinase dimer interface is closely similar to that observed in RAF homodimers (Figure 2C), which is not surprising considering that all interacting residues are identically conserved between BRAF and CRAF (Supplementary Figure 4D). As would be expected for an active dimer, both kinase domains exhibit an active “C-helix-in”, “DFG-in” conformation with their regulatory and catalytic spines aligned.^30^ Type I RAF inhibitor GDC-0879 was included in the protein preparation and there is density for the compound in the active sites of both BRAF and CRAF (Figure 2D).

The MEK1-bound complex is similar, but with MEK1 associated with the upward-facing BRAF subunit (Figure 2B). The MEK1 and BRAF kinase domains interact in the face-to-face orientation as seen in prior structures. The overall conformation observed here is similar to that seen previously in a BRAF homodimer structure with a single MEK1 bound (PDB entry 6Q0T, RMSD 2.28 Å for 1295 aligned Cα residues, Supplementary Figure 4C). We note that MEK1 is not as well-resolved as the RAF and 14-3-3 portions of this complex, likely due to conformational heterogeneity. It is also possible that particles lacking MEK1 remain in this reconstruction, despite our efforts to remove them via heterogeneous refinement.

Interestingly, we observe density corresponding to the BRAF NtA motif in this structure – the SSDD sequence extends across the dimer interface to contact the CRAF kinase domain (Figure 3A). The mainchain carbonyl of Ser447 in BRAF is positioned to hydrogen bond with Lys399 in CRAF, and Asp448 in BRAF forms a salt bridge with Arg398 in CRAF. The final residue in the BRAF SSDD motif, Asp449, forms an intra-protomer salt bridge with Arg506 (Figure 3A). This interaction is asymmetric; the position of the BRAF SSDD segment sterically precludes a similar interaction of the corresponding region of CRAF (we are using a CRAF mutant in which the native SSYY sequence is substituted with SSDD). The BRAF NtA motif is known to be constitutively phosphorylated on Ser446/447 to approximately 50% occupancy, but the observed interaction does not appear to be phosphorylation-dependent. The sidechains of both serine residues are solvent exposed, and we expect that the observed interaction could occur irrespective of their phosphorylation state.

**Figure 3.**
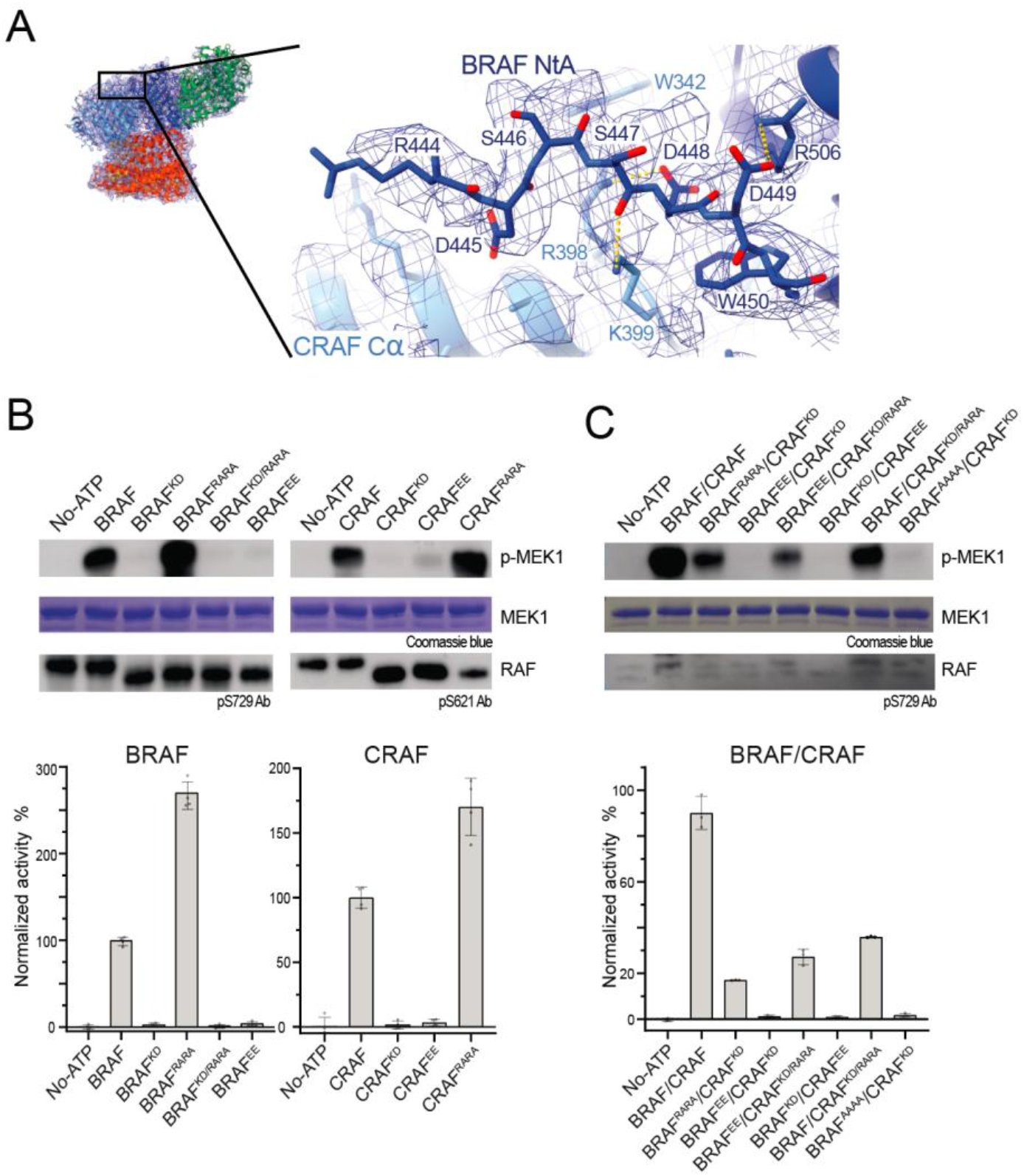
Structure-function analysis of the NtA motif in BRAF and CRAF. A, Cryo-EM map and interactions of the BRAF NtA motif in the BRAF/CRAF^FERA^/MEK1/14-3-3 structure. Ser447 and Asp448 in the BRAF NtA motif interact with Arg398 and Lys399 in the RKTR motif in the CRAF αC-helix (yellow dashed lines). Asp449 in the BRAF NtA motif is positioned to form an intra-protomer salt bridge with Arg506. B, Activity assays (MEK phosphorylation) with active or kinase-dead BRAF and CRAF homodimers with mutations in the NtA and RKTR motifs. RARA denotes replacement of the SSDD sequence of the NtA motif with RARA, EE denotes replacement of the arginine and lysine in the RKTR motif with glutamic acid residues, and KD denotes a kinase-dead mutant (D576N in BRAF and D468N in CRAF). BRAF and CRAF have an SSDD NtA motif except when marked as RARA. MEK1 phosphorylation was measured using the TR-FRET kinase assay and mean activity and standard deviation are shown in the bar graph (n=4). Products of one representative experiment were analyzed by SDS-PAGE with western blotting for phospho-MEK and Coomassie-staining of the MEK1 substrate and western blotting for RAF pSer621 or pSer729 as loading controls. C, As in B, but with BRAF/CRAF heterodimers, as indicated.

Because the density for the SSDD segment is somewhat weak and we do not observe density for this motif in our heterodimer structure lacking MEK, we assessed this interaction with molecular dynamics. A 1ms simulation showed a persistent interaction between the sidechains of Asp448 in BRAF and Arg398 in CRAF, despite considerable variability in the backbone conformation of the BRAF SSDD segment overall (Figure 4A,B). Owing to this overall flexibility, the intra-protomer salt bridge between Asp449 and Arg506 in BRAF was variably observed over the course of the simulation and the BRAF Ser447-CRAF Lys399 interaction was not stable; rather Lys399 more often interacted with the carbonyl of Ser446 in BRAF (Figure 4B).

**Figure 4.**
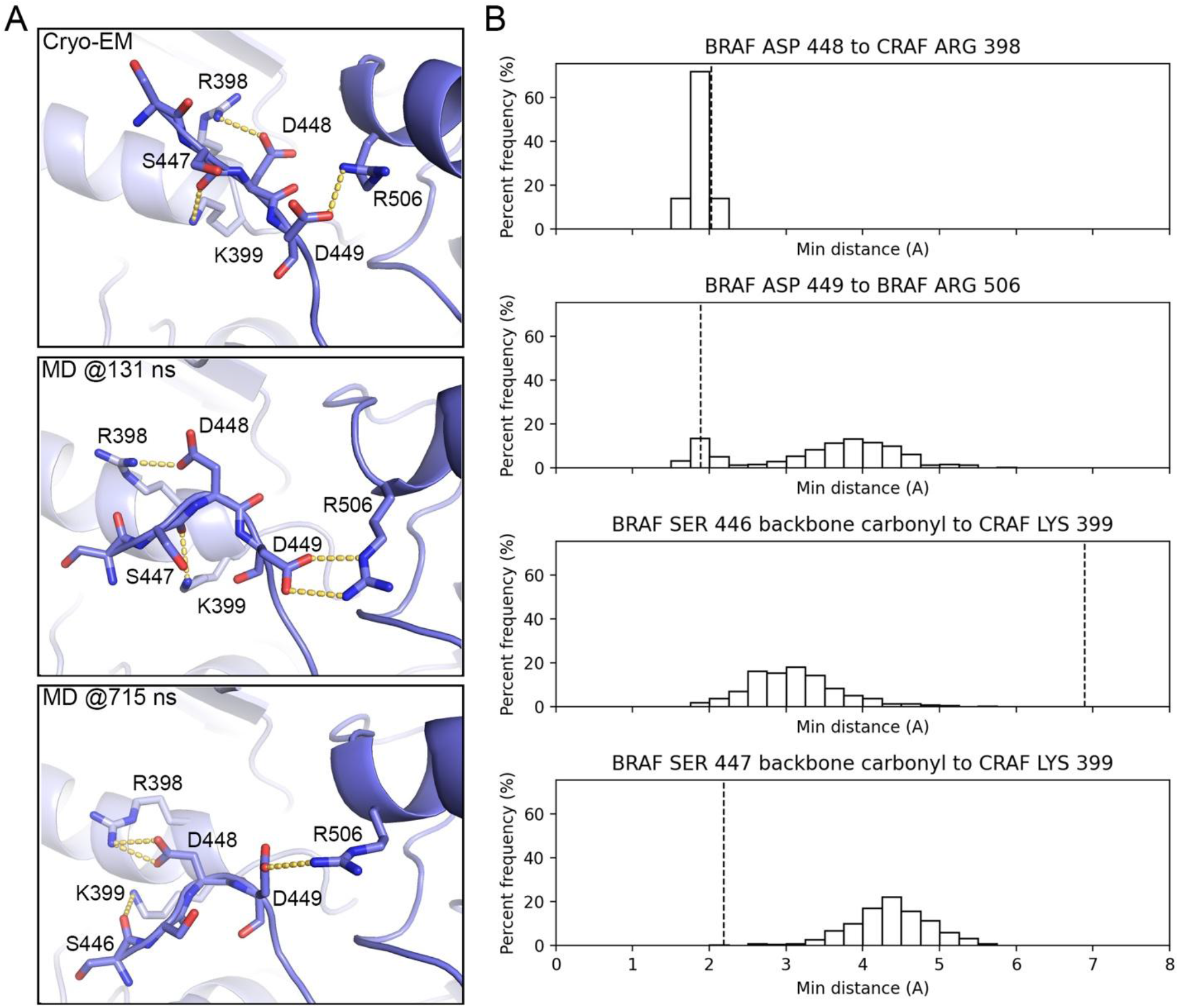
Evaluation of the SSDD interaction using molecular dynamics (MD). A, Interactions of the BRAF SSDD segment in the cryo-EM structure (top panel) and representative frames of a 1 µs MD simulation (lower two panels). BRAF is shown in dark blue and CRAF in light blue. Potential hydrogen bond interactions are shown as dashed yellow lines. B, Histograms showing frequency of occurrence of selected interactions of SSDD segment during a 1 µs MD simulation. Minimum distance (Å) is measured as closest approach of any of the atoms (including hydrogen atoms) in the indicated groups during 1 ns intervals of the simulation. Vertical dashed lines indicate distances in the cryo-EM structure (after addition of hydrogen atoms and initial minimization).

Prior work has highlighted the need for negative charge in the NtA motif of BRAF and CRAF, and Arg398 and Lys399 in CRAF are part of the conserved RKTR motif, which has also been shown to be crucial for RAF activity.^23,31-33^. Furthermore, these segments have been proposed to interact in *trans* to promote or stabilize RAF homodimers or heterodimers. ^27,34^ In light of these studies and our equivocal structural findings, we further probed the role of this interaction with mutagenesis of the SSDD and RKTR motifs, as described in the following section.

### Negative charge is not required in the NtA motif of BRAF or CRAF

Inspired by previous structure-function studies of transactivation in kinase dimers^27,35^, we designed a charge-swap experiment in which we replaced the negatively charged SSDD segment in BRAF or CRAF with the positively charged “RARA” and the RK portion of the interacting RKTR motif with the negatively charged “EE”. These variants were incorporated into BRAF and CRAF with or without a kinase-dead mutation (denoted BRAF^KD^ and CRAF^KD^) with the goal of ascertaining whether the activating effect of the NtA interaction was exerted in *cis* or in *trans* (Supplementary Figure 5). We co-expressed the resulting variants, including BRAF/CRAF heterodimer pairs, and tested the activity of the purified dimers/heterodimers in an *in vitro* MEK phosphorylation assay.

The lack of soluble expression or differing expression levels in several pairs of constructs confounded execution and interpretation of the original experiment as designed (Supplementary Figure 5). However, an unexpected finding emerged from the constructs that we were able to express and purify – replacement of the SSDD sequence with RARA yielded highly active BRAF and CRAF homodimers and heterodimers (Figure 3B, C). Indeed, BRAF^RARA^ and CRAF^RARA^ homodimers were more active than BRAF and CRAF homodimers bearing the SSDD sequence (BRAF^SSDD^ and CRAF^SSDD^, Figure 3B, lower panels). By contrast, the EE substitution in the RKTR motif largely abolished the activity of both BRAF and CRAF homodimers (Figure 3B, C). In BRAF/CRAF heterodimers, both the BRAF^RARA^/CRAF^KD^ and BRAF^SSDD^/CRAF^KD/RARA^ pairs were active, albeit less so than the BRAF^SSDD^/CRAF^SSDD^ heterodimer which contains two active kinase protomers (Figure 3C). A quadruple alanine substitution in BRAF yielded an inactive heterodimer with kinase-dead CRAF (BRAF^AAAA^/CRAF^KD^). Interestingly, co-expression of BRAF^EE^ with kinase-dead CRAF^KD/RARA^ yielded an active dimer, whereas co-expression with kinase-dead CRAF^KD/SSDD^ did not (Figure 3C). Considering these results and our structural observations, we conclude that while the NtA motif can interact across the dimer interface (Figure 3A), this interaction not required for RAF activity. Replacement of the acidic SSDD sequence with the basic RARA sequence yields highly active RAF homodimers and heterodimers.

## Discussion

Our biochemical and structural studies of the BRAF/CRAF/14-3-3 heterodimer complex reveals properties that are closely similar to those of BRAF and CRAF homodimers. This is not surprising considering the relatively high degree of sequence conservation between these RAF isoforms (~80% identity in the kinase domain). While the heterodimer was modestly more active than either homodimer, observed Michaelis constants for both ATP and MEK1 were essentially unchanged as compared with the homodimers. Our prior studies of the drug sensitivity of RAF isoforms showed that they can differ substantially between BRAF and CRAF, in particular for type II inhibitors which as a class are more potent inhibitors of CRAF. Here, the IC_50_ values we measured for the heterodimer were typically intermediate between those of the homodimers. For type II inhibitors, we further noticed a clear trend for inhibition to more closely parallel that observed for the less-sensitive BRAF homodimers (Supplementary Fig. 1, Table 1).

The cryo-EM structure of the BRAF/CRAF/MEK1/14-3-3 complex reveals an asymmetric interaction in which the BRAF NtA motif extends across the kinase dimer interface to contact the RKTR motif of the CRAF protomer. Although this interaction is consistent with prior biochemical observations implicating both the NtA and RKTR motifs as critical determinants of RAF catalytic activity^23,31-33^, we observed it only in the MEK1-bound structure and not in the heterodimer structure lacking MEK1. Furthermore, MD simulations revealed that while the Asp448–Arg398 contact was persistent, the broader NtA backbone was flexible and other interactions seen in the cryo-EM structure were only variably maintained. Together, these findings suggest that the NtA–RKTR contact is not a rigid, obligate feature of the active RAF heterodimer.

In further probing the NtA-RKTR contact with mutagenesis, we found that replacement of the acidic SSDD sequence with the basic RARA sequence yielded highly active BRAF and CRAF homodimers and heterodimers (Figure 3B,C). This result was unanticipated given the longstanding view that negative charge in the NtA region is required for RAF activation^1^, a view supported by the constitutive phosphorylation of the serine residues in BRAF’s NtA motif and the strong activating effect of serine and tyrosine phosphorylation (or acidic substitutions) in the ARAF and CRAF NtA motifs.^20,22,25,26,36^ The RARA substitution demonstrates that negative charge is not required, and notwithstanding the possibility of a neomorphic mechanism of activation, it argues against a simple model in which the native NtA drives activity by engaging the basic RKTR motif in trans. While our structure shows one way in which such an interaction can occur, the totality of the evidence argues against a requirement for stereospecific recognition of the NtA motif in an active RAF dimer or heterodimer. First, aspartic acid or glutamic acid substitutions in the CRAF NtA are not reasonable stereochemical “mimics” of phosphotyrosine and could not be expected to make comparable interactions at the dimer interface, yet they are highly activating. Second, the NtA motif is largely disordered in the many available structures of BRAF and CRAF homodimers and is variably ordered in the BRAF/CRAF heterodimer structures we report here. In further support of the variable nature of NtA interactions, a recent BRAF crystal structure reveals an asymmetric homodimer in which the Asp448 and Asp449 in SSDD motif of one protomer make interactions across the dimer interface^37^, but they are not analogous to those we observe here in the BRAF/CRAF heterodimer, even though the residues involved are conserved. Finally, as noted above, our mutagenesis experiments show that even the basic RARA substitution is activating in both BRAF and CRAF.

In a recent study of the structure of autoinhibited CRAF, we proposed that phosphorylation or mutation of the NtA segment could alter CRAF dimerization and activity via an effect on the stability of the inactive, monomeric state of the kinase^13^. This was based on comparison of structures of inactive CRAF with a native SSYY versus SSDD-substituted NtA motif. Although the NtA motif was not resolved in either structure, the inactive, αC-helix-out conformation was much better defined in the structure with the SSYY motif, and we speculated that phosphorylation or negative charge in the NtA motif could destabilize the monomer leading to an increased propensity to dimerize. The NtA segment is immediately adjacent to a conserved tryptophan residue (Trp342 in CRAF, Trp450 in BRAF) that inserts between the αC-helix and β4-strand in the N-lobe. In Src-family kinases, the corresponding residue plays a well-established role in maintaining the inactive, αC-helix-out conformation of the kinase^38,39^, and mutation of this residue in BRAF has been shown to impair activation^27^. Our present mutagenesis results are consistent with a model in which a charged NtA sequence — whether acidic or basic — could destabilize this tryptophan-anchored inactive conformation by increasing the dynamics of the adjacent backbone, thereby lowering the energetic barrier to formation of the αC-helix-in dimeric state. Future solution NMR studies focused on this tryptophan in RAFs with a phosphorylated versus non-phosphorylated NtA motif could provide the most direct test of this model. Importantly, we do not discount the importance of the interactions of the BRAF SSDD motif observed in our BRAF/CRAF/MEK1/14-3-3 heterodimer structure as they may also contribute to dimerization or heterodimerization, if in a more ephemeral manner.Finally, we note that while this NtA interaction is asymmetric, both BRAF and CRAF kinase domains protomers adopt an active configuration in the 14-3-3-bound heterodimer and our findings do not support a “transactivation” model in which the NtA segment is proposed to promote activation of the opposing subunit in the dimer^27,37^.

## Methods

### Preparation of RAF kinase homodimers and heterodimers

Recombinant baculovirus expressing human BRAF and CRAF kinase constructs indicated below were prepared using baculoviral transfer vector pAc8. For protein production Sf9 insect cells were infected with high-titer stocks of baculovirus expressing either BRAF or CRAF (for preparation of homodimers) or were co-infected with both BRAF-and CRAF-expressing baculovirus (for preparation of heterodimers) using standard procedures.^40^ For selected preparations as indicated, MEK1 was also co-expressed by co-infection with baculovirus expressing full-length MEK1 S218A/S222A (MEK^SASA^) with an N-terminal 6xHis tag.

| RAF | Residues | Tag | Designation | Mutation |
| --- | --- | --- | --- | --- |
| BRAF | 435-766 | N-terminal<br>Strep | BRAF | - |
|  |  |  | BRAF<sup>KD</sup> | D576N |
|  |  |  | BRAF<sup>RARA</sup> | 446-RARA-449 |
|  |  |  | BRAF<sup>KD/RARA</sup> | D576N, 446-RARA-449 |
|  |  |  | BRAF<sup>EE</sup> | 506-EE-507 |
|  |  |  | BRAF<sup>AAAA</sup> | 446-AAAA-449 |
| CRAF | 308-648,<br>Y340D/Y341D | N-terminal<br>6xHis | CRAF or CRAF<sup>SSDD</sup> | (all CRAF constructs except CRAF<sup>SSYY</sup> contain Y340D/Y341D) |
|  |  |  | CRAF<sup>SSYY</sup> | Y340/Y341 |
|  |  |  | CRAF<sup>KD</sup> | D468N |
|  |  |  | CRAF<sup>RARA</sup> | 338-RARA-341 |
|  |  |  | CRAF<sup>KD/RARA</sup> | D468N/<br>338-RARA-341 |
|  |  |  | CRAF<sup>EE</sup> | 398-EE-399 |
|  |  |  | CRAF<sup>AAAA</sup> | 338-AAAA-341 |
|  |  |  | CRAF<sup>FERA</sup> | F559E/R563A |

Baculovirus-infected cells were harvested and lysed in lysis buffer (25 mM Tris-HCl pH 8.0, 0.15M NaCl, 5mM MgCl_2_, 1mM TCEP, 1 μM ATP-γ-S, 10% glycerol) with protease inhibitor cocktail (Thermo Fisher Scientific) and RAF dimers were purified by affinity chromatography on HisTrap HP (Cytiva) and/or StrepTrap HP (Cytiva) as follows. BRAF homodimers were purified using StrepTrap HP in binding buffer (100mM Tris-HCl pH 8.0, 0.15M NaCl, 5mM MgCl_2_, 1mM TCEP, 1 μM ATP-γ-S) and elution buffer (binding buffer with 5mM desthiobiotin) followed by size-exclusion chromatography on Superdex 200 increase 10/300GL (Cytiva) in SEC buffer (25mM Tris-HCl pH 8.0, 0.15M NaCl, 5mM MgCl_2_, 1mM TCEP, 1 μM ATP-γ-S). CRAF homodimers were purified using HisTrap HP affinity chromatography in binding buffer (25 mM Tris-HCl pH 8.0, 0.15M NaCl, 5mM MgCl_2_, 1mM TCEP, 1 μM ATP-γ-S) and elution buffer (biding buffer containing 500mM Imidazole) followed by size-exclusion chromatography (Superdex 200 increase 10/300GL) in SEC buffer. BRAF/CRAF heterodimers were prepared by serial affinity chromatography on HisTrap HP and StrepTrap HP followed by size-exclusion chromatography using the buffers and conditions above. For purifications with co-expressed MEK1^SASA^, MEK inhibitor GDC-0623 (2 μM) was added to SEC buffer. For purification of the BRAF/CRAF^FERA^/MEK1 complex, SEC buffer was supplemented with both GDC-0623 (2 μM) and GDC-0879 (2 μM). All RAF dimers and heterodimers co-purified with insect-cell-derived 14-3-3ε and 14-3-3ζ as previously reported.^40^

### RAF kinase assays and inhibitor titrations

HTRF kinase assays were performed with the indicated RAF homo- or hetero-dimers and biotinylated MEK1 as substrate essentially as previously described.^28,40^ Briefly, standard conditions included 1 nM of the indicated RAF kinase preparation and 250 nM biotinylated MEK1. Reactions were initiated upon the addition of ATP (200 μM final), incubated for 30 minutes, then quenched upon the addition of detection buffer supplemented with 62.5 nM XL665 and 0.5 nM of a europium-coupled anti-phospho MEK1/2 antibody. After a 40 minute incubation, the FRET signal ratio 665nm/620nm was measured using a PHERAstar microplate reader (BMG Labtech). For comparisons of homo-versus heterodimer activity and CRAF versus CRAF^FERA^ activity, the indicated range of concentrations of RAF enzyme was employed under the standard assay conditions. For experiments assessing the effect of NtA and RKTR motif mutations on the activity of RAF homo- and hetero-dimers, enzyme concentrations were as follows: 15nM for BRAF homodimers, 10nM for CRAF homodimers, and 20nM for BRAF/CRAF heterodimers.

For K_m[ATP]_ and K_m[MEK]_ determinations, ATP and biotinylated MEK1 concentrations were titrated as indicated. The ATP titrations were fit in GraphPad PRISM with a substrate inhibition model to determine K_m[ATP]_ and K_i[ATP]_ values. The MEK titrations were fit in GraphPad PRISM with the Michaelis-Menten model to determine K_m[MEK]_ values.

For inhibitor titrations, all RAF dimers were assayed at a final concentration of 1 nM using the standard conditions above, and were pre-incubated with inhibitors for 40 minutes before a 30-minute reaction with ATP and a 40-minute detection period. The resulting concentration-response curves were fit with a four-parameter inhibition model in GraphPad PRISM to determine IC_50_ and Hill Slopes (nH) for each inhibitor and enzyme combination, as reported in Table 1.

For kinase activity assays using western blot detection, reactions were carried out using standard conditions as for HTRF but reactions were stopped by addition of 1x SDS sample buffer and heating to 95°C in a heat block. After SDS-PAGE, MEK1 phosphorylation was detected by western blotting with antibodies for MEK1/2 (Cell Signaling Technology, #9122) or phospho-MEK1/2 (Cell Signaling Technology, #9121).

### MEK1 pulldown assay

To test binding of CRAF and CRAF^FERA^ to MEK1, Strep-tagged MEK1 (5ug) was incubated with 20uL Strep-Tactin beads (iba) in binding buffer (25 mM Tris-HCl pH 8.0, 0.15M NaCl, 5mM MgCl_2_, 1mM TCEP, 1 μM ATP-γ-S) for 30 min at 4°C. Next, 6xHis-tagged CRAF or CRAF^FERA^ was added to the MEK1-bound beads and incubated for 60 min at 4°C with agitation. His_6_-tagged BRAF^V600E^ was used as an additional positive control. After three washes, the beads were eluted with SDS-PAGE sample buffer and bound proteins were analyzed by SDS-PAGE and western blotting with an anti-His_5_ antibody.

### Cryo-EM imaging and single particle reconstruction

The BRAF/CRAF^FERA^/MEK1^SASA^/14-3-3 complex in SEC buffer with GDC-0623 and GDC-0879 (2μM each) was diluted to 0.6 mg/mL and applied to glow-discharged holey gold grids (Quantifoil UltraAUFoil 1.2/1.3, 300 mesh). Grids were blotted at 95% humidity at 10°C for 3.5 sec and vitrified with a Leica EM-GP. A total of 9,676 micrograph movies were collected on a Titan Krios at 300 kV with Gatan Quantum Image Filter with K2 direct detector at nominal magnification of 105,000 x in counting mode with a pixel size of 0.825 Å/pixel. 63 frames were recorded per movie (3.5 sec exposure time) with defocus varying from −0.8 to −2.2 μm with a total dose of 62.5 e^-^/Å^2^(see Supplementary Table 1).

The movies were motion-corrected by Patch Motion Correction and CTF was estimated by Patch CTF Estimation in CryoSPARC.^41^ Particle picking with TOPAZ^42^ yielded 2,588,280 particles from 8,236 curated micrographs (Supplementary Figure 3). Particles were then re-extracted in RELION^43^ from micrographs that were pre-processed by MotionCor2 and CTFFIND4.1 within RELION. Initial rounds of 2D classification and 3D *ab initio* reconstruction allowed identification of two distinct complexes, one containing MEK1 (BRAF/CRAF^FERA^/MEK1/14-3-3, 831,958 particles) and one without MEK1 (BRAF/CRAF^FERA^/14-3-3, 1,343,109 particles). For the BRAF/CRAF^FERA^/MEK1/14-3-3 complex, further rounds of ab initio reconstruction, 3D classification, and 3D auto-refinement led to selection of 171,208 particles which after CTF-Refinement and Bayesian polishing led to 3.9 Å reconstruction for the MEK1-containing complex using Fourier shell correlation (FSC) 0.143 threshold criterion.

For the BRAF/CRAF^FERA^/14-3-3 complex, the 1,343,109 particles were used for *ab initio* reconstruction followed by heterogeneous refinement allowing selection of 528,370 particles (Supplementary Figure 3). A final round of ab initio reconstruction in CryoSPARC led to selection of 139,194 particles which were imported to RELION for CTF-Refinement and Bayesian polishing, which resulted in the final 3.3 Å reconstruction.

### Cryo-EM model building and refinement

Atomic models were created by manual docking of relevant domains from PDB entries 6NYB and 6PP9 to the cryo-EM density using Chimera X. ^44^ Automated refitting was carried out with ModelAngelo.^45^ The resulting models were then manually inspected and refit using COOT ^46^ and final models were refined against the density maps using Phenix’s Real-space refinement with secondary structure restraints and Ramachandran restraints. In the BRAF/CRAF^FERA^/14-3-3 complex, chain A residues 603-610, chain B residues 496-508, 517-522, and 563-566 were modeled based on reference PDB files due to weak cryo-EM density. In the BRAF/CRAF^FERA^/MEK1/14-3-3 complex, chain A residues 603-612 and 657-658, chain B residues 498-509, 519-522, and 561-566, and chain C residues 63-67, 84-86, 155-159, and 366-378, were modeled based on reference PDB files due weak cryo-EM density.

### Molecular Dynamics Simulations

The system preparation and molecular dynamics simulations described here were carried out as outlined below using previously reported protocols.^18^ Initial structures were derived from the BRAF/CRAF^FERA^/MEK1/14-3-3 heterodimer structure and the system was prepared using the Protein Preparation Wizard in Schrödinger Suite, including hydrogen atom addition, H-bond network optimization, and the determination of protonation states appropriate for physiological pH. To mitigate potential steric overlaps, a restrained energy minimization was applied.^47^ Ligands were extracted from the system for parameterization, and ligand parameters were generated using the AMBER ff19sb force field.^48^ Small molecule structures in SDF format were processed using AmberTools suite to derive parameters.^49^ AM1-BCC charges were calculated using Antechamber, with the GAFF2 force field parameters assigned via parmchk2. To integrate these with the OpenMM engine, parameters were serialized into XML format.^50^ Protein-ligand complexes were prepared by merging the extracted ligands and then solvating the systems in a TIP3P water box with a padding of 1.2 nm buffer.^51^ Counter-ions were added to neutralize the system at a physiological ionic strength of 0.1 M. Energy minimization was performed using a convergence-based approach with a force tolerance of 1.0 kJ/mol/nm and an energy tolerance of 0.1 kJ/mol over 50 steps. A total of 10000 steps was used for this process.

The system was equilibrated using a dual-stage thermalization strategy. The initial phase involved a linear temperature ramp from 50 K to 310 K over a 2.5 ns duration. During the first 1.25 ns of this ramp, harmonic positional restraints on the protein backbone atoms were gradually phased out, decaying from an initial force constant of 10.0 to 0.0 kJ/mol/nm^2^. This was followed by a 1.0 ns isothermal hold at 310 K to ensure kinetic stabilization.

For the MD simulation, production-level sampling for each system was conducted for a total of 1000 ns within the NPT ensemble (310 K, 1.0 bar). We utilized a Langevin integrator with a collision frequency of 1.0 ps^‒1^ for temperature regulation and a Monte Carlo barostat (25-step update frequency) for pressure control. To maximize computational throughput, Hydrogen Mass Repartitioning (HMR) was employed— reassigning hydrogen masses to 4.0 Da—which permitted an integration timestep of 4.0 fs.^52^ The production run was performed for 1000 ns. Long-range electrostatic forces were calculated using the Particle Mesh Ewald (PME) method under periodic boundary conditions. A 1.0 nm cutoff was applied to all non-bonded interactions. Structural integrity was maintained by constraining all covalent bonds via the SHAKE algorithm.^53,54^ Convergence and system stability were verified through the post-hoc monitoring of total energy and Root Mean Square Deviation (RMSD). All trajectory processing and structural metrics were computed using the MDTraj library.^55^

A separate post-processing protocol was implemented to quantify ligand–protein interaction energies from the production trajectories using OpenMM. For each system, the final equilibrated topology and coordinates and corresponding trajectory were analyzed with MDTraj to identify interactions of the BRAF NtA motif.^55^ A custom OpenMM system was then constructed using a force field stack consisting of protein.ff19SB, phosaa19SB, and ligand-specific parameters generated via GAFF2 XML parameter files. Reported distances were then averaged over the last 500 frames to obtain mean and standard deviation values.

All stages, from force-field assignment to simulation execution, were automated using custom Python workflows (available on GitHub). Workflow scripts are available at https://github.com/dgazgalis/Workflows-Python-Scripts.

## Acknowledgments

This work was supported in part by grants R35CA242461 (MJE) and P50CA165962 (MJE) from the National Cancer Institute of the U.S. National Institutes of Health and by the Pediatric Low-grade Astrocytoma Program at the Dana-Farber Cancer Institute.

## Declaration of generative AI and AI-assisted technologies in the writing process

During the preparation of this work the authors used Claude to edit the manuscript. After using this tool/service, the authors reviewed and edited the content as needed and take full responsibility for the content of the published article.

## References

1. Lavoie, H., and Therrien, M. (2015). Regulation of RAF protein kinases in ERK signalling. Nature reviews. Molecular cell biology 16, 281–298. 10.1038/nrm3979.

2. Jeon, H., Tkacik, E., and Eck, M.J. (2024). Signaling from RAS to RAF: The Molecules and Their Mechanisms. Annu Rev Biochem 93, 289–316. 10.1146/annurev-biochem-052521-040754.

3. Simanshu, D.K., and Morrison, D.K. (2022). A Structure is Worth a Thousand Words: New Insights for RAS and RAF Regulation. Cancer Discov 12, 899–912. 10.1158/2159-8290.Cd-21-1494.

4. Davies, H., Bignell, G.R., Cox, C., Stephens, P., Edkins, S., Clegg, S., Teague, J., Woffendin, H., Garnett, M.J., Bottomley, W., et al. (2002). Mutations of the BRAF gene in human cancer. Nature 417, 949–954. 10.1038/nature00766.

5. Poulikakos, P.I., Persaud, Y., Janakiraman, M., Kong, X., Ng, C., Moriceau, G., Shi, H., Atefi, M., Titz, B., Gabay, M.T., et al. (2011). RAF inhibitor resistance is mediated by dimerization of aberrantly spliced BRAF(V600E). Nature 480, 387–390. 10.1038/nature10662.

6. Jones, D.T.W., Bandopadhayay, P., and Jabado, N. (2019). The Power of Human Cancer Genetics as Revealed by Low-Grade Gliomas. Annu Rev Genet 53, 483–503. 10.1146/annurev-genet-120417-031642.

7. Imielinski, M., Greulich, H., Kaplan, B., Araujo, L., Amann, J., Horn, L., Schiller, J.,Villalona-Calero, M.A., Meyerson, M., and Carbone, D.P. (2014). Oncogenic and sorafenib-sensitive ARAF mutations in lung adenocarcinoma. The Journal of clinical investigation 124, 1582–1586. 10.1172/JCI72763.

8. Noeparast, A., Giron, P., Noor, A., Bahadur Shahi, R., De Brakeleer, S., Eggermont, C., Vandenplas, H., Boeckx, B., Lambrechts, D., De Greve, J., and Teugels, E. (2019). CRAF mutations in lung cancer can be oncogenic and predict sensitivity to combined type II RAF and MEK inhibition. Oncogene 38, 5933–5941. 10.1038/s41388-019-0866-7.

9. Karoulia, Z., Gavathiotis, E., and Poulikakos, P.I. (2017). New perspectives for targeting RAF kinase in human cancer. Nature reviews. Cancer 17, 676–691. 10.1038/nrc.2017.79.

10. Hymowitz, S.G., and Malek, S. (2018). Targeting the MAPK Pathway in RAS Mutant Cancers. Cold Spring Harb Perspect Med 8. 10.1101/cshperspect.a031492.

11. Park, E., Rawson, S., Li, K., Kim, B.W., Ficarro, S.B., Pino, G.G., Sharif, H., Marto, J.A., Jeon, H., and Eck, M.J. (2019). Architecture of autoinhibited and active BRAF-MEK1-14-3-3 complexes. Nature 575, 545–550. 10.1038/s41586-019-1660-y.

12. Martinez Fiesco, J.A., Durrant, D.E., Morrison, D.K., and Zhang, P. (2022). Structural insights into the BRAF monomer-to-dimer transition mediated by RAS binding. Nat Commun 13, 486. 10.1038/s41467-022-28084-3.

13. Jang, D.M., Boxer, K., Ha, B.H., Tkacik, E., Levitz, T., Rawson, S., Metivier, R.J., Schmoker, A., Jeon, H., and Eck, M.J. (2025). Cryo-EM structures of CRAF/MEK1/14-3-3 complexes in autoinhibited and open-monomer states reveal features of RAF regulation. Nat Commun 16, 8150. 10.1038/s41467-025-63227-2.

14. Kondo, Y., Ognjenovic, J., Banerjee, S., Karandur, D., Merk, A., Kulhanek, K., Wong, K., Roose, J.P., Subramaniam, S., and Kuriyan, J. (2019). Cryo-EM structure of a dimeric B-Raf:14-3-3 complex reveals asymmetry in the active sites of B-Raf kinases. Science 366, 109–115. 10.1126/science.aay0543.

15. Liau, N.P.D., Wendorff, T.J., Quinn, J.G., Steffek, M., Phung, W., Liu, P., Tang, J., Irudayanathan, F.J., Izadi, S., Shaw, A.S., et al. (2020). Negative regulation of RAF kinase activity by ATP is overcome by 14-3-3-induced dimerization. Nature structural & molecular biology 27, 134–141. 10.1038/s41594-019-0365-0.

16. Dedden, D., Nitsche, J., Schneider, E.V., Thomsen, M., Schwarz, D., Leuthner, B., and Gradler, U. (2024). Cryo-EM Structures of CRAF(2)/14-3-3(2) and CRAF(2)/14-3-3(2)/MEK1(2) Complexes. J Mol Biol 436, 168483. 10.1016/j.jmb.2024.168483.

17. Ryan, M.B., Quade, B., Schenk, N., Fang, Z., Zingg, M., Cohen, S.E., Swalm, B.M., Li, C., Ozen, A., Ye, C., et al. (2024). The Pan-RAF-MEK Nondegrading Molecular Glue NST-628 Is a Potent and Brain-Penetrant Inhibitor of the RAS-MAPK Pathway with Activity across Diverse RAS- and RAF-Driven Cancers. Cancer Discov 14, 1190–1205. 10.1158/2159-8290.CD-24-0139.

18. Chakraborty, S., Jiang, J., Camallonga, J.V., Gazgalis, D., Gero, T.W., Tavares, M.T., Dittemore, G., Tkacik, E., Jang, D.M., Ha, B.H., et al. (2025). RAF isoform selectivity of MEK inhibitors and rational design of a covalent ARAF-MEK inhibitor. bioRxiv. 10.64898/2025.12.15.693503.

19. Terrell, E.M., and Morrison, D.K. (2019). Ras-Mediated Activation of the Raf Family Kinases. Cold Spring Harb Perspect Med 9. 10.1101/cshperspect.a033746.

20. Marais, R., Light, Y., Paterson, H.F., and Marshall, C.J. (1995). Ras recruits Raf-1 to the plasma membrane for activation by tyrosine phosphorylation. EMBO J 14, 3136–3145.

21. Jelinek, T., Dent, P., Sturgill, T.W., and Weber, M.J. (1996). Ras-induced activation of Raf-1 is dependent on tyrosine phosphorylation. Mol Cell Biol 16, 1027–1034. 10.1128/MCB.16.3.1027.

22. Mason, C.S., Springer, C.J., Cooper, R.G., Superti-Furga, G., Marshall, C.J., and Marais, R. (1999). Serine and tyrosine phosphorylations cooperate in Raf-1, but not B-Raf activation. EMBO J 18, 2137–2148. 10.1093/emboj/18.8.2137.

23. Emuss, V., Garnett, M., Mason, C., and Marais, R. (2005). Mutations of C-RAF are rare in human cancer because C-RAF has a low basal kinase activity compared with B-RAF. Cancer research 65, 9719–9726. 10.1158/0008-5472.CAN-05-1683.

24. Cook, F.A., and Cook, S.J. (2021). Inhibition of RAF dimers: it takes two to tango. Biochem Soc Trans 49, 237–251. 10.1042/BST20200485.

25. Fabian, J.R., Daar, I.O., and Morrison, D.K. (1993). Critical tyrosine residues regulate the enzymatic and biological activity of Raf-1 kinase. Mol Cell Biol 13, 7170–7179. 10.1128/mcb.13.11.7170-7179.1993.

26. Diaz, B., Barnard, D., Filson, A., MacDonald, S., King, A., and Marshall, M. (1997). Phosphorylation of Raf-1 serine 338-serine 339 is an essential regulatory event for Ras-dependent activation and biological signaling. Mol Cell Biol 17, 4509–4516. 10.1128/MCB.17.8.4509.

27. Hu, J., Stites, E.C., Yu, H., Germino, E.A., Meharena, H.S., Stork, P.J.S., Kornev, A.P., Taylor, S.S., and Shaw, A.S. (2013). Allosteric activation of functionally asymmetric RAF kinase dimers. Cell 154, 1036–1046. 10.1016/j.cell.2013.07.046.

28. Tkacik, E., Jang, D.M., Boxer, K., Ha, B.H., and Eck, M.J. (2025). Characterization and inhibitor sensitivity of ARAF, BRAF, and CRAF kinases. J Biol Chem 301, 110800. 10.1016/j.jbc.2025.110800.

29. Garnett, M.J., Rana, S., Paterson, H., Barford, D., and Marais, R. (2005). Wild-type and mutant B-RAF activate C-RAF through distinct mechanisms involving heterodimerization. Mol Cell 20, 963–969. 10.1016/j.molcel.2005.10.022.

30. Taylor, S.S., and Kornev, A.P. (2011). Protein kinases: evolution of dynamic regulatory proteins. Trends Biochem Sci 36, 65–77. 10.1016/j.tibs.2010.09.006.

31. Baljuls, A., Mueller, T., Drexler, H.C., Hekman, M., and Rapp, U.R. (2007). Unique Nregion determines low basal activity and limited inducibility of A-RAF kinase: the role of N-region in the evolutionary divergence of RAF kinase function in vertebrates. J Biol Chem 282, 26575–26590. 10.1074/jbc.M702429200.

32. Baljuls, A., Mahr, R., Schwarzenau, I., Muller, T., Polzien, L., Hekman, M., and Rapp, U.R. (2011). Single substitution within the RKTR motif impairs kinase activity but promotes dimerization of RAF kinase. J Biol Chem 286, 16491–16503. 10.1074/jbc.M110.194167.

33. Takahashi, M., Li, Y., Dillon, T.J., Kariya, Y., and Stork, P.J.S. (2017). Phosphorylation of the C-Raf N Region Promotes Raf Dimerization. Mol Cell Biol 37. 10.1128/MCB.00132-17.

34. Jambrina, P.G., Rauch, N., Pilkington, R., Rybakova, K., Nguyen, L.K., Kholodenko, B.N., Buchete, N.V., Kolch, W., and Rosta, E. (2016). Phosphorylation of RAF Kinase Dimers Drives Conformational Changes that Facilitate Transactivation. Angew Chem Int Ed Engl 55, 983–986. 10.1002/anie.201509272.

35. Zhang, X., Gureasko, J., Shen, K., Cole, P.A., and Kuriyan, J. (2006). An allosteric mechanism for activation of the kinase domain of epidermal growth factor receptor. Cell 125, 1137–1149. 10.1016/j.cell.2006.05.013.

36. Brummer, T., Martin, P., Herzog, S., Misawa, Y., Daly, R.J., and Reth, M. (2006). Functional analysis of the regulatory requirements of B-Raf and the B-Raf(V600E) oncoprotein. Oncogene 25, 6262–6276. 10.1038/sj.onc.1209640.

37. Kondo, Y., Notbohm, J., Navas Camacho, I., Nagy-Davidescu, G., Mason, T., Mühle, J., Standfuss, J., and Perica, T. (2025). Mechanism of MEK1 phosphorylation by the N-terminal acidic motif-mediated asymmetric BRAF dimer. bioRxiv, 2025.2009.2026.678760. 10.1101/2025.09.26.678760.

38. LaFevre-Bernt, M., Sicheri, F., Pico, A., Porter, M., Kuriyan, J., and Miller, W.T. (1998). Intramolecular regulatory interactions in the Src family kinase Hck probed by mutagenesis of a conserved tryptophan residue. J Biol Chem 273, 32129–32134. 10.1074/jbc.273.48.32129.

39. Joseph, R.E., Xie, Q., and Andreotti, A.H. (2010). Identification of an allosteric signaling network within Tec family kinases. J Mol Biol 403, 231–242. 10.1016/j.jmb.2010.08.035.

40. Tkacik, E., Li, K., Gonzalez-Del Pino, G., Ha, B.H., Vinals, J., Park, E., Beyett, T.S., and Eck, M.J. (2023). Structure and RAF family kinase isoform selectivity of type II RAF inhibitors tovorafenib and naporafenib. J Biol Chem 299, 104634. 10.1016/j.jbc.2023.104634.

41. Punjani, A., Rubinstein, J.L., Fleet, D.J., and Brubaker, M.A. (2017). cryoSPARC: algorithms for rapid unsupervised cryo-EM structure determination. Nat Methods 14, 290–296. 10.1038/nmeth.4169.

42. Bepler, T., Morin, A., Rapp, M., Brasch, J., Shapiro, L., Noble, A.J., and Berger, B. (2019). Positive-unlabeled convolutional neural networks for particle picking in cryo-electron micrographs. Nat Methods 16, 1153–1160. 10.1038/s41592-019-0575-8.

43. Kucukelbir, A., Sigworth, F.J., and Tagare, H.D. (2014). Quantifying the local resolution of cryo-EM density maps. Nat Methods 11, 63–65. 10.1038/nmeth.2727.

44. Meng, E.C., Goddard, T.D., Pettersen, E.F., Couch, G.S., Pearson, Z.J., Morris, J.H., and Ferrin, T.E. (2023). UCSF ChimeraX: Tools for structure building and analysis. Protein Sci 32, e4792. 10.1002/pro.4792.

45. Jamali, K., Kall, L., Zhang, R., Brown, A., Kimanius, D., and Scheres, S.H.W. (2024). Automated model building and protein identification in cryo-EM maps. Nature 628, 450–457. 10.1038/s41586-024-07215-4.

46. Emsley, P., Lohkamp, B., Scott, W.G., and Cowtan, K. (2010). Features and development of Coot. Acta Crystallogr D Biol Crystallogr 66, 486–501. 10.1107/S0907444910007493.

47. Sastry, G.M., Adzhigirey, M., Day, T., Annabhimoju, R., and Sherman, W. (2013). Protein and ligand preparation: parameters, protocols, and influence on virtual screening enrichments. J Comput Aided Mol Des 27, 221–234. 10.1007/s10822-013-9644-8.

48. Bradbury, M.W., Stump, D., Guarnieri, F., and Berk, P.D. (2011). Molecular modeling and functional confirmation of a predicted fatty acid binding site of mitochondrial aspartate aminotransferase. J Mol Biol 412, 412–422. 10.1016/j.jmb.2011.07.034.

49. Case, D.A., Aktulga, H.M., Belfon, K., Cerutti, D.S., Cisneros, G.A., Cruzeiro, V.W.D., Forouzesh, N., Giese, T.J., Gotz, A.W., Gohlke, H., et al. (2023). AmberTools. J Chem Inf Model 63, 6183–6191. 10.1021/acs.jcim.3c01153.

50. Shirts, M.R., Klein, C., Swails, J.M., Yin, J., Gilson, M.K., Mobley, D.L., Case, D.A., and Zhong, E.D. (2017). Lessons learned from comparing molecular dynamics engines on the SAMPL5 dataset. J Comput Aided Mol Des 31, 147–161. 10.1007/s10822-016-9977-1.

51. Cisneros, G.A., Wikfeldt, K.T., Ojamae, L., Lu, J., Xu, Y., Torabifard, H., Bartok, A.P., Csanyi, G., Molinero, V., and Paesani, F. (2016). Modeling Molecular Interactions in Water: From Pairwise to Many-Body Potential Energy Functions. Chem Rev 116, 7501–7528. 10.1021/acs.chemrev.5b00644.

52. Hopkins, C.W., Le Grand, S., Walker, R.C., and Roitberg, A.E. (2015). Long-Time-Step Molecular Dynamics through Hydrogen Mass Repartitioning. J Chem Theory Comput 11, 1864–1874. 10.1021/ct5010406.

53. Eastman, P., Swails, J., Chodera, J.D., McGibbon, R.T., Zhao, Y., Beauchamp, K.A., Wang, L.P., Simmonett, A.C., Harrigan, M.P., Stern, C.D., et al. (2017). OpenMM 7: Rapid development of high performance algorithms for molecular dynamics. PLoS Comput Biol 13, e1005659. 10.1371/journal.pcbi.1005659.

54. Eastman, P., Galvelis, R., Pelaez, R.P., Abreu, C.R.A., Farr, S.E., Gallicchio, E., Gorenko, A., Henry, M.M., Hu, F., Huang, J., et al. (2024). OpenMM 8: Molecular Dynamics Simulation with Machine Learning Potentials. J Phys Chem B 128, 109–116. 10.1021/acs.jpcb.3c06662.

55. McGibbon, R.T., Beauchamp, K.A., Harrigan, M.P., Klein, C., Swails, J.M., Hernandez, C.X., Schwantes, C.R., Wang, L.P., Lane, T.J., and Pande, V.S. (2015). MDTraj: A Modern Open Library for the Analysis of Molecular Dynamics Trajectories. Biophys J 109, 1528–1532. 10.1016/j.bpj.2015.08.015.

