## Supplemental data for "Cryo-EM structure and biochemical characterization of a BRAF/CRAF heterodimer: Negative charge in the NtA motif is not required for RAF activation"

This file includes:

Supplemental Table 1

Supplemental Figures 1 to 5

**Supplementary Table 1. Cryo-EM data collection, refinement and validation statistics**

| **Data collection and processing** | | |
|---|---|---|
| Magnification | 105,000x | |
| Voltage (kV) | 300 | |
| Electron exposure (e/Å$^2$) | 62.5 | |
| Defocus range (μm) | - 0.8 to -2.2 | |
| Pixel size | 0.825 | |
| Symmetry imposed | C1 | |
| Number of micrographs | 9,676 | |
| **Refinement** | BRAF/CRAF$^{FERA}$/14-3-3 complex | BRAF/CRAF$^{FERA}$/MEK1$^{SASA}$/14-3-3 complex |
| PDB | 13DU | 13DV |
| EMD | EMD-77011 | EMD-77012 |
| Final particle images (no.) | 139,194 | 171,208 |
| Map resolution (Å) | | |
| 0.143 FSC threshold | 3.3 | 3.9 |
| Initial model used | 6NYB | 6NYB, 6PP9 |
| Model composition | | |
| Chains | 4 | 5 |
| Non-hydrogen atoms | 8,301 | 10,399 |
| Protein residues | 1,028 | 1,295 |
| Ligands | 29L (2) | 29L (2), LCJ (1) |
| B factors (Å$^2$) | | |
| Proteins | 65.5 | 112.0 |
| Ligands | 45.2 | 97.6 |
| R.m.s deviations | | |
| Bond length (Å) | 0.002 | 0.003 |
| Bond angles (°) | 0.457 | 0.570 |
| Validation | | |
| MolProbity score | 2.83 | 3.41 |
| Ramachandran plot | | |
| Favored (%) | 94.85 | 90.81 |
| Outliers (%) | 0 | 0 |
| Model vs. Data | | |
| CC (mask) | 0.76 | 0.64 |
| CC (box) | 0.71 | 0.68 |
| CC (peaks) | 0.64 | 0.52 |
| CC (volume) | 0.75 | 0.64 |
| Mean CC for ligands | 0.76 | 0.63 |

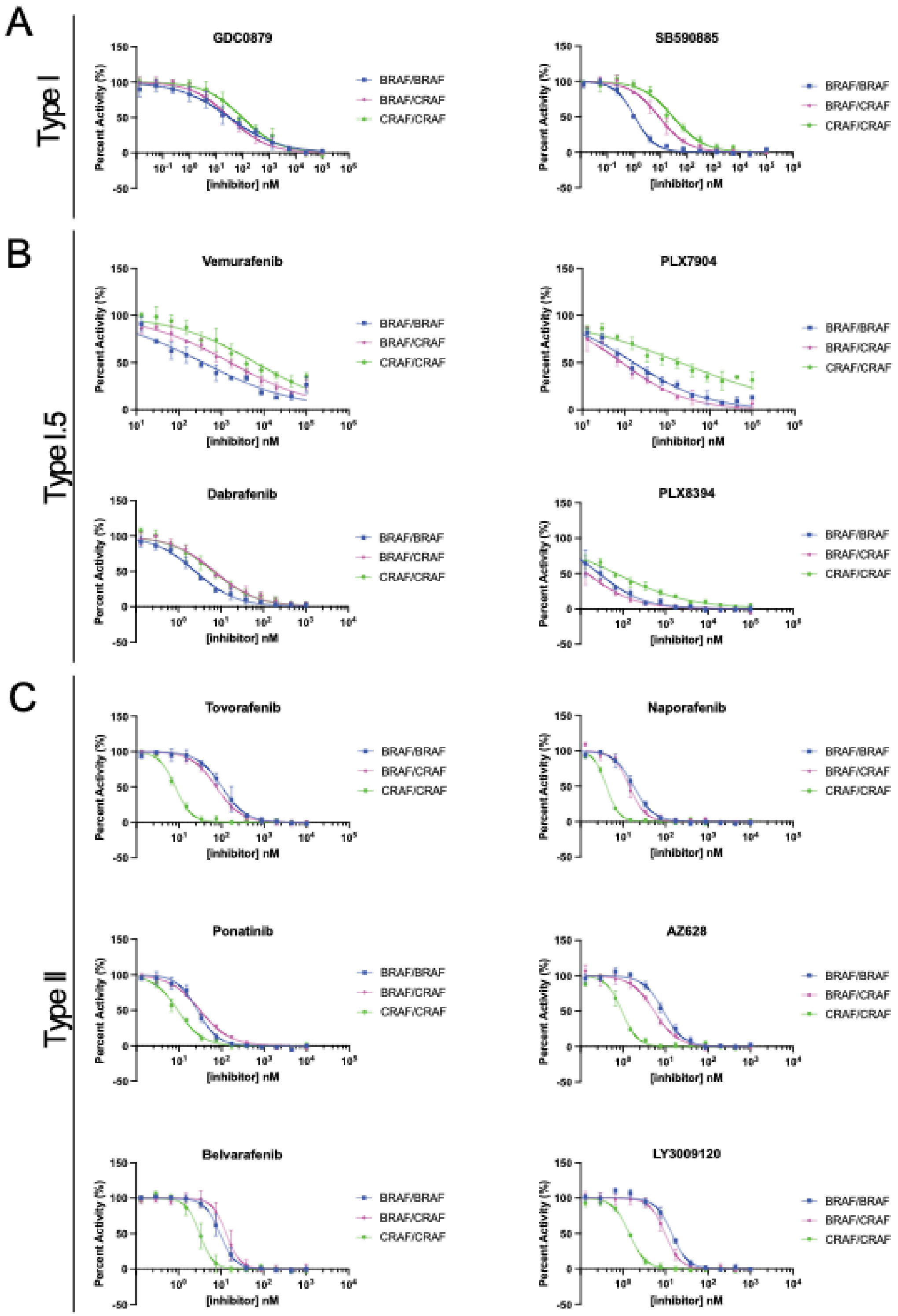


**Supplementary Figure 1. Profiling of RAF inhibitors across BRAF and CRAF homodimers and heterodimers.** Representative concentration-response curves used to determine $IC_{50}$ values provided in Table 1 of the main text. Data are plotted as mean ± standard deviation from one independent experiment performed in triplicate (n≥3).

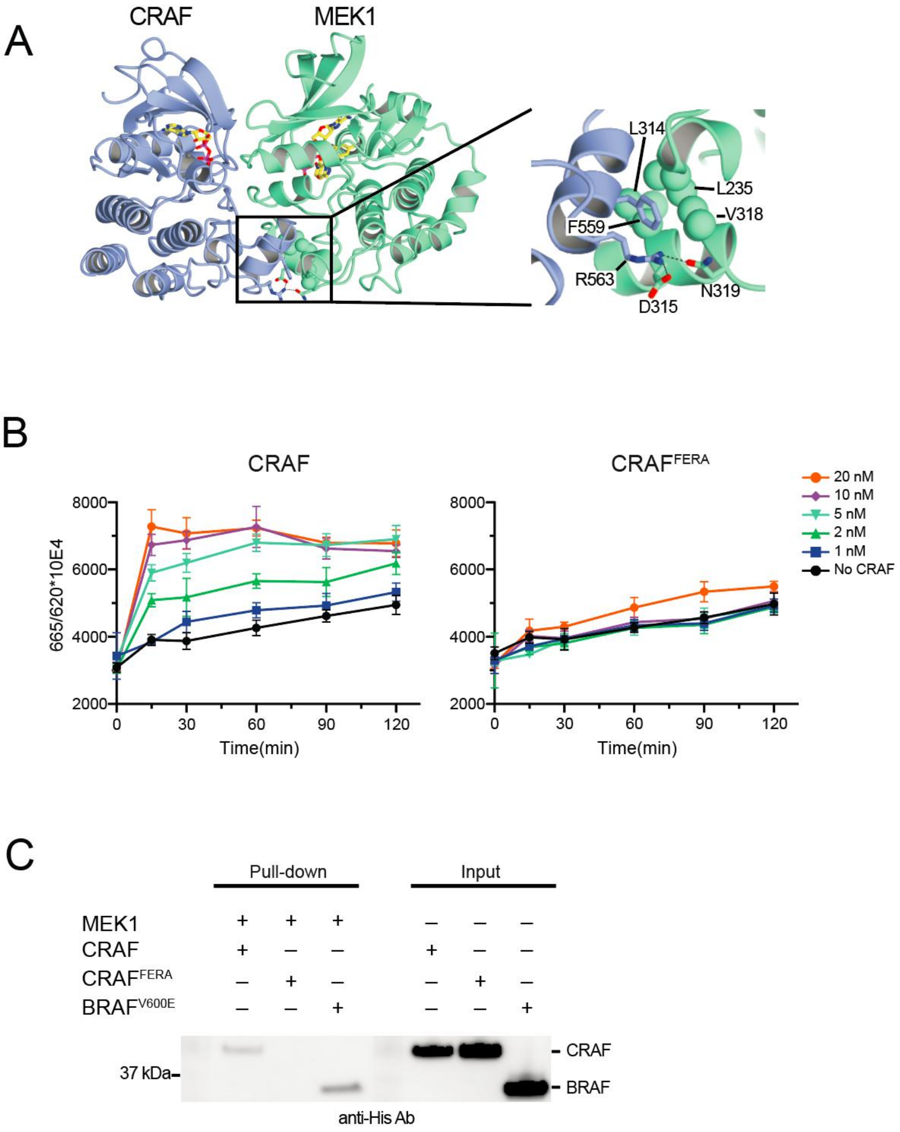


**Supplementary Figure 2. Design and evaluation of a CRAF variant that does not bind MEK.** A, Guided by our prior cryo-EM structure of CRAF and MEK1 (PDB: 9MMQ), we introduced the F559E/R563A double mutation in the $\alpha$G-helix of CRAF to prevent its association with MEK1. We refer to the resulting construct as $CRAF^{FERA}$ and it also contains the SSDD substitution of the NtA motif. CRAF is shown in light blue and MEK1 in green, and the MEK1 residues interacting with F559 and R563 are indicated. B, The activity of CRAF and $CRAF^{FERA}$ was measured in a TR-FRET assay, varying the concentration of CRAF and the incubation time. As expected, $CRAF^{FERA}$ was markedly impaired in its ability to phosphorylate MEK1. Values plotted are mean ± standard deviation (n=3) C, Pull-down assay testing binding of CRAF, $CRAF^{FERA}$, or $BRAF^{V600E}$ to bead-immobilized MEK1. MEK1 with an N-terminal strep tag was first immobilized on strep beads at 4°C. The MEK1-bound beads were then incubated with $His_6$-tagged CRAF, $CRAF^{FERA}$, or $BRAF^{V600E}$ for 1 hour at 4°C. After three washing steps, the beads were eluted with SDS-PAGE sample buffer and bound proteins were analyzed by SDS-PAGE and blotting with an anti-$His_5$ antibody. The 37 kDa molecular weight marker is indicated. The three lanes on the right show 10% of the input of each RAF protein.

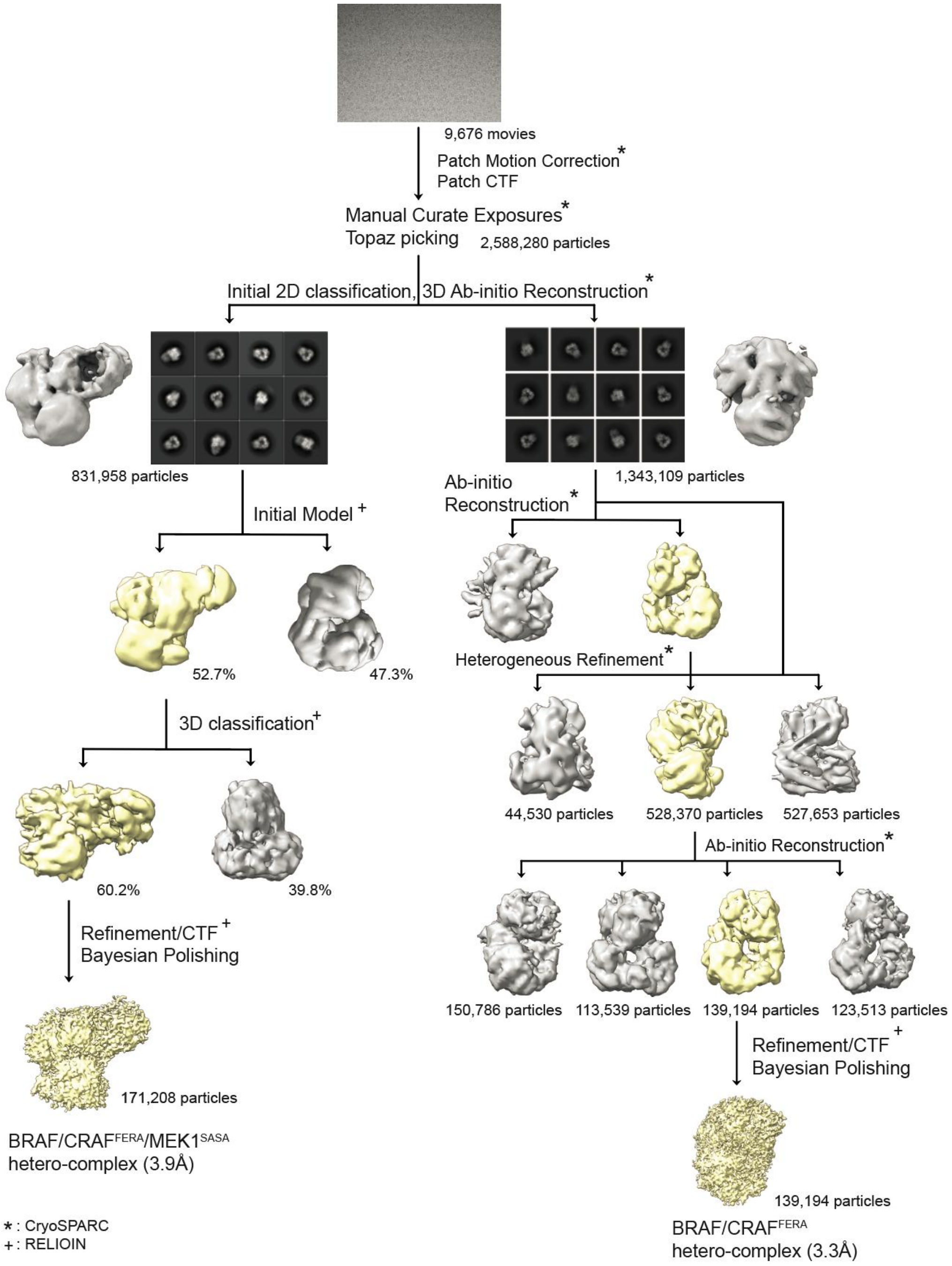


**Supplementary Figure 3. Cryo-EM image processing workflow.** A total of 9,676 movie micrographs were recorded using a Thermo Fisher Titan Krios cryo-TEM equipped with a K2 direct electron detector and Gatan Quantum Image filter. Patch motion correction and patch CTF correction were performed using CryoSPARC, followed by manual curation of the exposures and particle picking with Topaz. This resulted in the selection of 2,588,280 particles, which were then subjected to 2D classification and initial model generation using Ab-initio methods. The selected particles were transferred to RELION and processed through CTF refinement and Bayesian polishing to produce the final map. "*" indicates processes performed in CryoSPARC, and "+" indicates processes processed using RELION.

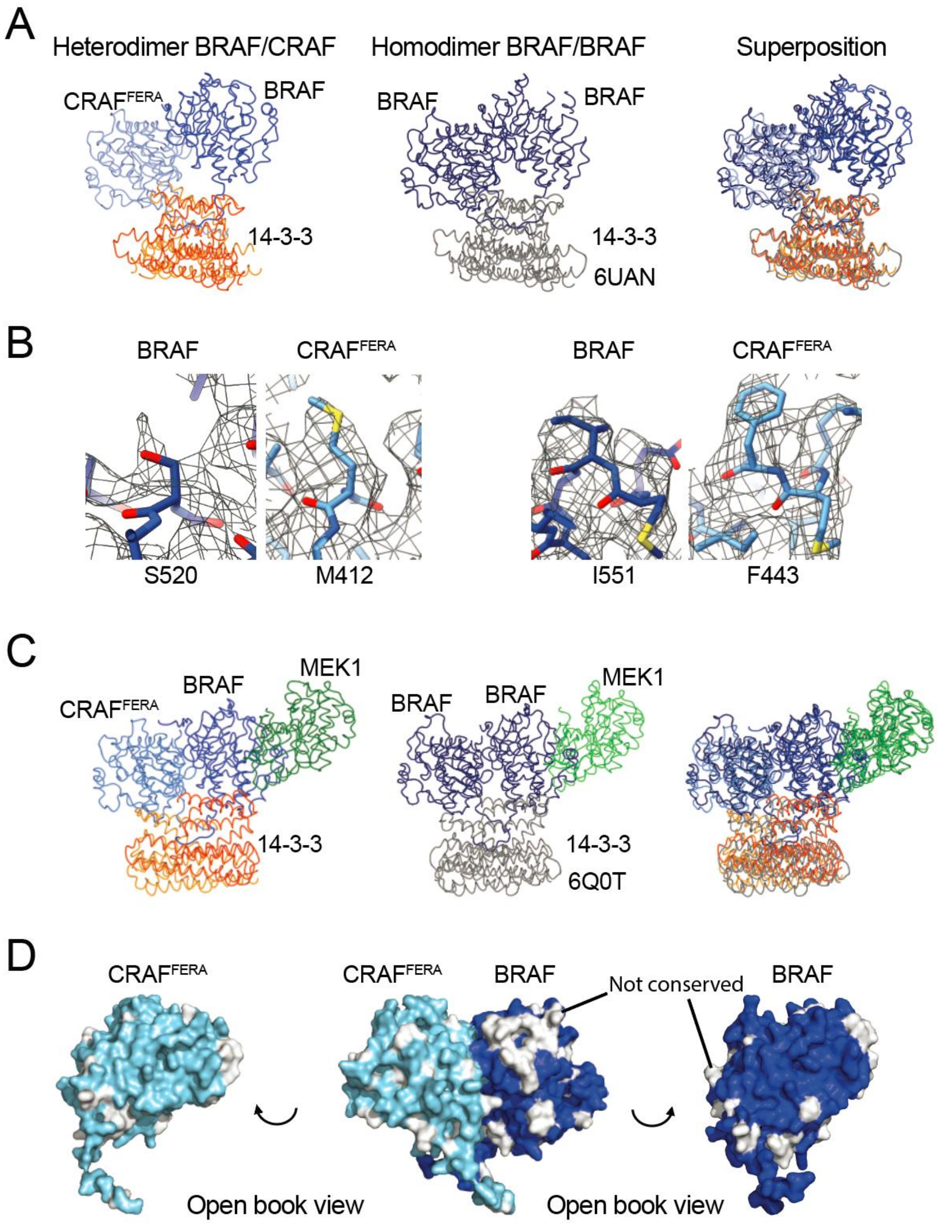


**Supplementary Figure 4. Additional structural analysis of the BRAF/CRAF heterodimer.** A, Comparison of the BRAF/CRAF/14-3-3 heterodimer structure with that of the previously determined BRAF homodimer (PDB: 6UAN). Superposition of the two cryo-EM structures is shown on the far right (r.m.s.d. = 1.65Å for 1007 aligned Cα atoms). B, Cryo-EM density for selected sidechains that differ in size in BRAF versus CRAF. Differences in sidechain density for these and other residues guided assignment of the corresponding protomers in the heterodimer. C, Comparison of the MEK-bound BRAF/CRAF/14-3-3 heterodimer structure with that of the previously determined BRAF/MEK1/14-3-3 homodimer structure with a single MEK1 bound (PDB: 6Q0T). Superposition of the two cryo-EM structures is shown on the far right (r.m.s.d. = 1.65Å for 1007 aligned Cα atoms). D, "Open-book" view of the BRAF/CRAF dimer interface. Residues that are not identically conserved between BRAF and CRAF are shown in white on the surface models of BRAF (blue) and CRAF (light blue). Note that the interacting surfaces revealed in the open book views on the far left (for CRAF) and far right (for BRAF) are entirely conserved.

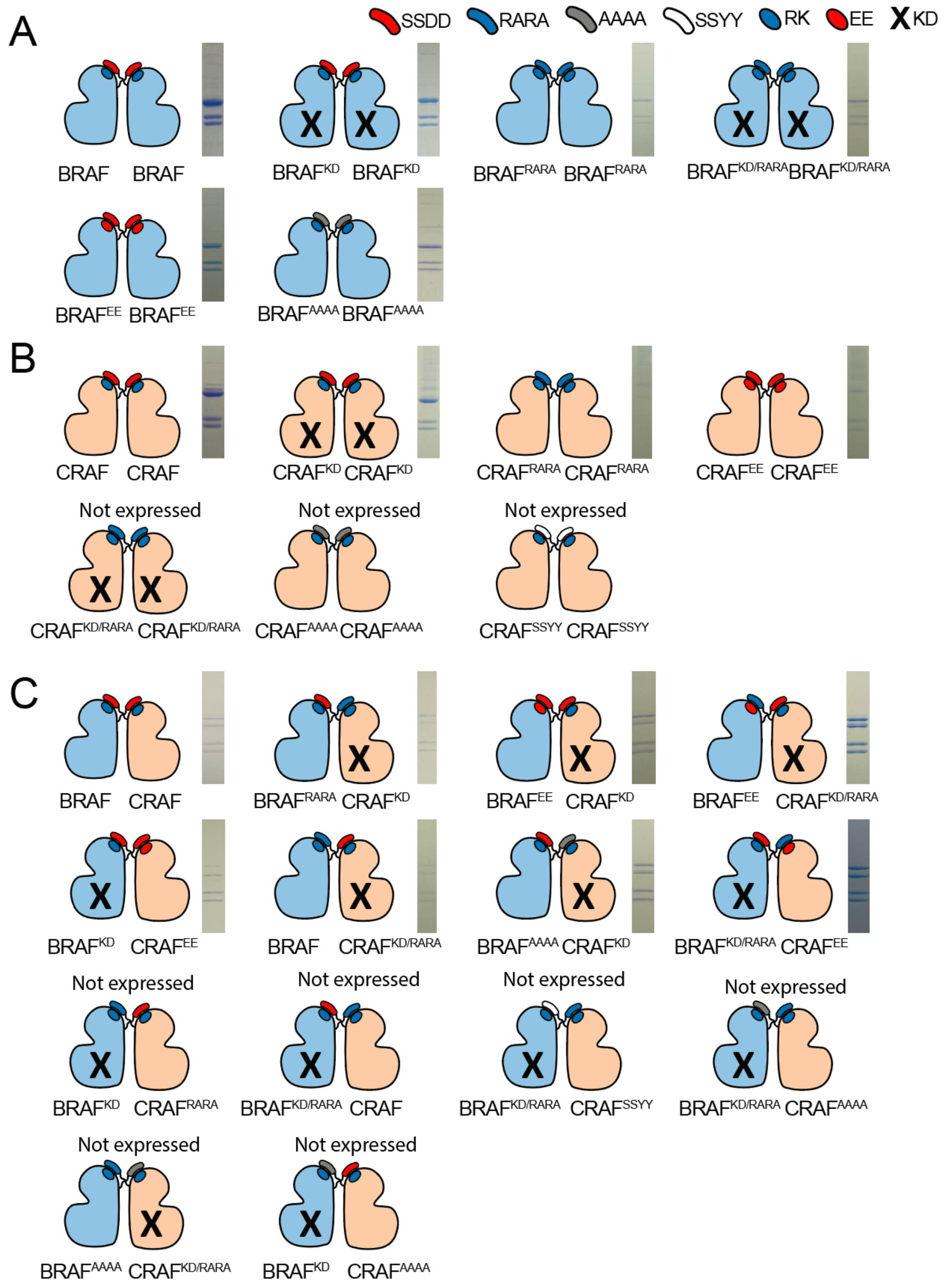


**Supplementary Figure 5. Design of a structure-function experiment to test the role of the NtA/RKTR motif interaction in RAF homodimers and heterodimers.** The key at the top of the figure identifies substitutions of the NtA and RKTR motifs in the context of (A) BRAF homodimers, (B) CRAF homodimers, or (C) BRAF/CRAF heterodimers. The NtA substitutions indicate the four-residue sequence of the NtA motif and the RKTR substitutions indicate the first two residues of this motif (RK or its replacement with EE). The "X" mark indicates a kinase-dead (KD) mutation (D576N in BRAF, D468N in CRAF). The corresponding lane of a Coomassie-stained SDS-PAGE gel is provided for the variants that were successfully expressed and purified as 14-3-3-bound dimers from insect cells. Activity assays for selected complexes are shown in Figure 3 in the main text.
